# MIC-Drop-seq: Scalable single-cell phenotyping of mutant vertebrate embryos

**DOI:** 10.1101/2025.05.27.656468

**Authors:** Clayton M. Carey, Saba Parvez, Zachary J. Brandt, Brent W. Bisgrove, Christopher J. Yates, Randall T. Peterson, James A. Gagnon

## Abstract

Pooled perturbation screens can reveal cellular regulatory networks, yet scaling these techniques for large-scale screens in animals remains challenging. To address this, we developed MIC-Drop-seq, which combines high-throughput CRISPR gene disruption in zebrafish embryos with phenotyping by multiplexed single-cell RNAseq. In one MIC-Drop-seq experiment, we simultaneously identified changes in gene expression and cell abundance across 74 cell types resulting from loss of function of 50 transcription factors. These observations recapitulated many known phenotypes, while also uncovering novel functions in brain and mesoderm development. A key advantage of whole-animal screens is that they reveal how changes in one cell type affect the development of other cell types. Surprisingly, such cell-extrinsic phenotypes were abundant, indicating that transcription factors frequently exert effects beyond the cells where they are expressed to adjacent cells. We propose that MIC-Drop-seq will facilitate efforts to dissect the complete gene regulatory networks that guide animal development.

## Main Text

Much of our understanding of biology derives from studying how genetic perturbations manifest in organismal phenotypes (*1*). Forward mutagenic screens, for instance, provided a foundational knowledge of genetic networks underlying animal development (*2–4*). The recent advent of reverse genetic strategies employing CRISPR has opened up new possibilities for targeted screens that enable rapid linking of genotypes to phenotypes (*5*). Such targeted approaches, however, have been limited in scope by cumbersome manual delivery of perturbations to individual animals. We recently developed Multiplexed Intermixed CRISPR Droplets (MIC-Drop) as a technique for high-throughput generation of libraries of barcoded mutant zebrafish (*6*). In a MIC-Drop screen, a library of intermixed droplets is created and injected dropwise into zebrafish embryos at the single-cell stage to generate a pool of whole-animal mutants. Each droplet contains unique Cas9/guide RNA (gRNA) complexes targeting a single gene alongside a DNA barcode to facilitate rapid genotyping. We previously employed MIC-Drop to rapidly identify new regulators of cardiac development, demonstrating its power to expand the scale of reverse genetic screens in zebrafish (*7*).

Genetic screens of laboratory animals are often limited to easily observable phenotypes best suited for detection by simple visual assays. However, many developmental phenotypes are subtle or lack obvious morphological changes, rendering them invisible to conventional screening methods. Furthermore, many experimental measures of phenotype are qualitative or bespoke to specific systems, making them difficult to compare across experiments and laboratories. The advent of high-throughput transcriptional profiling, including single-cell RNA sequencing (scRNAseq), has provided a means to systematically characterize and quantify mutant phenotypes at the cellular and molecular levels. Single-cell methods have been used to map time-resolved cell atlases of zebrafish development, providing a baseline to compare perturbed states in a developing vertebrate (*8–10*). Recent high-resolution scRNAseq-based comparisons of mutant zebrafish embryos revealed previously unappreciated developmental phenotypes, even for well-characterized transcriptional regulators (*11*). Scaling these approaches for screening large numbers of mutants, however, remains logistically challenging and cost-prohibitive due to the individual engineering, housing, dissociation and separate sequencing of animals with different genotypes.

Approaches that employ multiplexed perturbation and pooled scRNAseq offer a path forward. Such methods, including Perturb-seq and CROP-seq, use indexing of CRISPR gRNAs from pooled cells to assign genotypes across hundreds of gene knockouts in a single experiment (*12–14*). These tools have allowed the identification of unique cell states and changes in gene regulatory networks caused by perturbing genes in cultured cells (*15–17*). Perturb-seq has been extended to organ-targeted *in vivo* systems, allowing the characterization of many knockout phenotypes within clonal populations in the brains of individual mice using viral vectors (*18*). Existing *in vivo* Perturb-seq approaches, however, fail to capture early developmental phenotypes that would manifest in full knockout animals, and can be confounded by interactions between clones with different genotypes in the same tissue. A whole-animal Perturb-seq approach, where each individual receives a single perturbation, is therefore needed to fully characterize the developmental function of genes *in vivo*.

Here we present MIC-Drop-seq, a technique that couples rapid, multiplexed CRISPR perturbation with scRNAseq in whole-animals. We demonstrate that MIC-Drop-seq enables effective genotype inference by gRNA capture in mutant cells and recapitulates known mutant phenotypes at the cellular level. Using MIC-Drop-seq for a screen of 50 transcription factors, we discover and validate roles for several transcription factors in brain and mesoderm development. We demonstrate that disruptions of transcription factors often cause pervasive yet gene-specific cell-extrinsic effects, indicating that transcription factors frequently alter cellular behaviors beyond the cells in which they are expressed and highlighting the unique advantages that come from performing Perturb-seq experiments in animals. Overall, our findings showcase the utility of MIC-Drop-seq for high-throughput screens of animal development, accelerating the mapping of genotypes to phenotypes across dozens of genes in parallel.

### MIC-Drop-seq enables multiplexed single-cell phenotyping of mutant zebrafish embryos

MIC-Drop-seq uses a simple workflow to create and pool animals with different mutant genotypes for multiplexed scRNAseq (**Fig. 1A**). As in MIC-Drop (*6,7*), sets of droplets containing Cas9/gRNA complexes targeting different genes are intermixed to form a library. Cohorts of zebrafish embryos are injected at the single-cell stage with a single droplet from the library, so that each embryo has a single gene mutated. Pooled mutant embryos are dissociated into a single-cell suspension of mixed genotypes and prepared for transcriptional profiling. To facilitate demultiplexing of mutant genotypes, MIC-Drop-seq uses direct capture and sequencing of gRNA. We used gRNAs with a modified scaffold that incorporates a capture sequence compatible with 10X Genomics feature barcoding (*13*) (**Fig. S1A)**. We validated that these gRNAs persist for days after injection (**Fig. S1B**) and were highly active for CRISPR mutagenesis in zebrafish embryos (**Fig. S1C**). This approach enables the unified capture of both mRNA and gRNA from each cell in pools derived from zebrafish embryos with mixed genotypes. MIC-Drop-seq therefore enables the multiplexed generation and single-cell transcriptomic profiling of many whole-animal mutants in a single experiment.

**Fig. 1:**
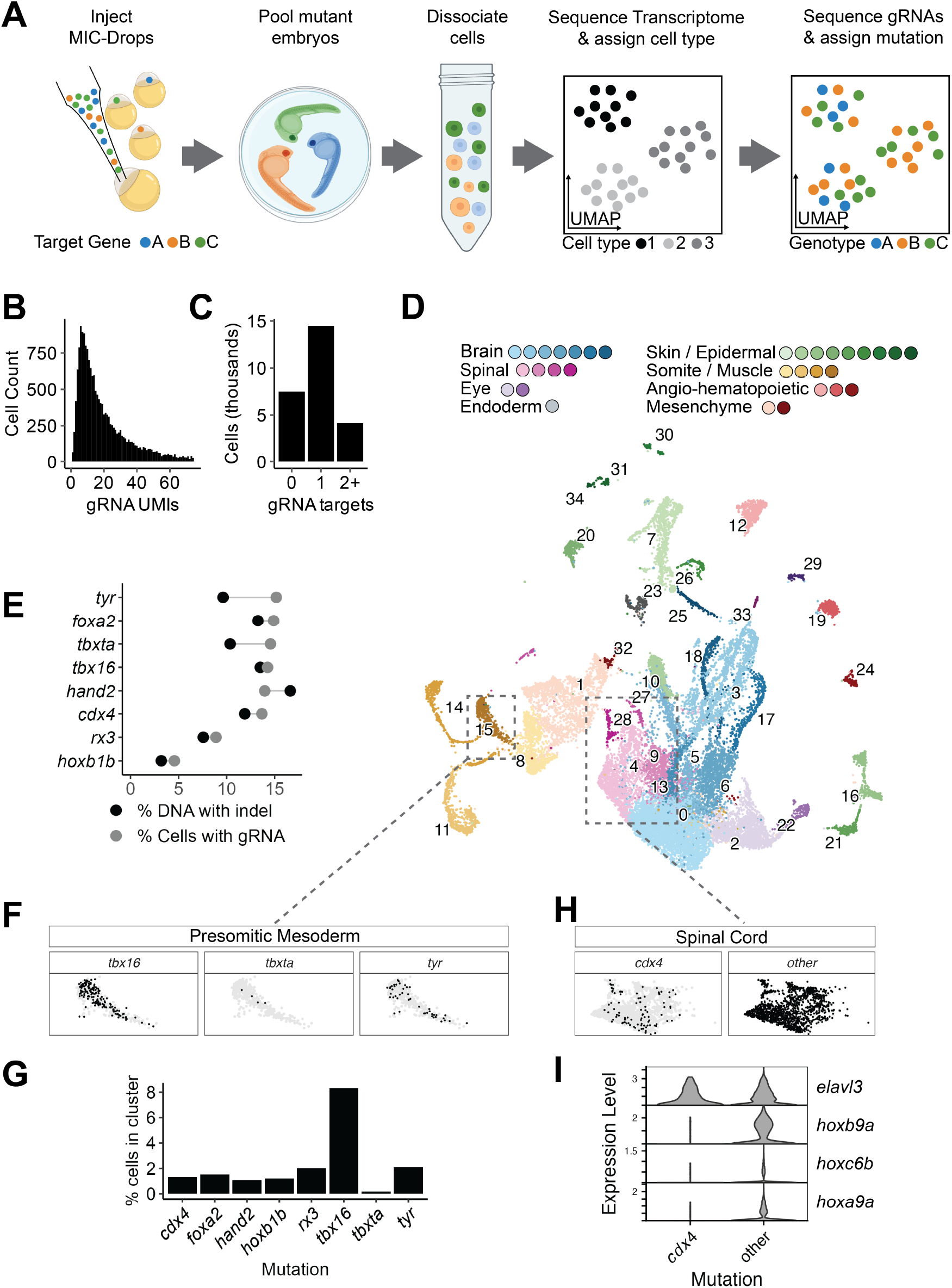
MIC-Drop-seq enables multiplexed single-cell phenotyping of mutant zebrafish embryos. (A) An overview of MIC-Drop-seq. (B) Histogram of unique guide RNA molecule counts sequenced in each cell with an assigned mutation. (C) Quantification of the number of cells in the dataset with no gRNAs detected (0), gRNA(s) targeting a single gene target (1) or gRNAs targeting multiple genes (2+). (D) UMAP embedding of 24 hpf zebrafish cells from a MIC-Drop-seq experiment targeting eight genes. (E) Comparison of observed DNA editing frequency and inferred genotypic composition of sequenced cells. The percentage of cells classified with the indicated genotype among all labeled cells was calculated (black) and compared to the observed indel frequency from bulk genomic DNA at each locus (grey) (F,H). Subsets of UMAP embeddings that highlight cells with the predicted genotype indicated (black) vs other cells (grey) in the (F) presomitic mesoderm and (H) spinal cord cluster(s). (G) Quantification of the percent of cells with the indicated genotype in the presomitic mesoderm cluster. (I) Gene expression violin plot comparing hox gene expression in *cdx4* mutant cells compared to all other cells in the spinal cord cluster. *elavl3* is a common marker of spinal cord cells whose expression is unchanged.

As a proof-of-concept, we performed a small MIC-Drop-seq screen targeting seven transcription factors with well-characterized mutant phenotypes (*tbx16, tbxta, cdx4, foxa2, hoxb1b, rx3*, and *hand2*), as well as a non-transcription factor control gene (*tyr*). A set of droplets was created for each gene target, with each droplet containing 4 unique gRNAs targeting the gene of interest and Cas9 protein, followed by intermixing to form a MIC-Drop library (**Table S1**). These droplets were used to inject a cohort of embryos at the single cell stage. A set of 40 individuals were then dissociated in a single pool at 24 hours post-fertilization (hpf) and processed together in a single super-loaded 10X channel. After sequencing the barcoded gRNA library, read thresholds were established to filter out ambient gRNA reads (see Methods). With these criteria, gRNAs were confidently detected in 71% of cells, with a median of 15 gRNA molecules recovered from each classified cell (**Fig. 1B**). Sequencing of gRNA targeting regions revealed that most cells had gRNAs targeting a single gene, while a smaller subset had gRNAs that targeted multiple genes (**Fig. 1C**). We reasoned that these cells represent doublets created by super-loading cells during microfluidic library preparation. Indeed, the proportion of cells with guides targeting multiple genes matched the expected doublet rate from super-loading. Computational doublet classification demonstrated that cells with gRNAs targeting multiple genes had characteristics of multiplet artifacts (**Fig. S2A**), and were more likely to be assigned as doublets (**Fig. S2B**)(*19*). We therefore considered these cells to be technical doublets and removed them from the dataset. The mutant genotype of the remaining 21,994 cells were inferred based on the sequence of the captured gRNAs, recovering a range of 661 to 2,198 cells per targeted gene (**Fig. S2A,B**). Dimensionality reduction and clustering yielded 35 cell clusters (**Fig. 1D**), which were annotated by label transfer from the Daniocell reference atlas (*8*) (**Fig.S4A-C**). Cluster annotations were manually verified by examining marker gene expression and represent all tissue types known to be present in the 24 hpf embryo (**Table S2**).

We determined the efficiency of CRISPR mutagenesis in our MIC-Drop-seq experiment using pooled amplicon sequencing of genomic DNA from the remaining dissociated cells. For each gene, we calculated the frequency of indels at the target cut sites and in experimental and wild type cells. We then compared editing frequency to the frequency of predicted mutant genotypes among cells in the dataset. The proportion of edited DNA and predicted mutant genotypes were highly correlated, indicating a high efficiency of mutagenesis (**Fig. 1E**), similar to previous MIC-Drop experiments (*7*).

We evaluated whether MIC-Drop-seq could recapitulate known developmental phenotypes. Two of the transcription factors we targeted have well-characterized roles in somitogenesis, each causing distinct changes in the abundance of specific mesodermal cell types. Embryos harboring mutations in *tbx16* have patterning defects in trunk somites and an accumulation of uncommitted presomitic mesoderm (*20*).

Conversely, mutations in *tbxta*, result in the failure to specify presomitic mesoderm and do not make posterior somites (*2*). We computed the proportion of cells in the presomitic mesoderm cluster for each genotype. Consistent with the known phenotypes, we observed a ∼4-fold enrichment of *tbx16* mutant cells and a ∼10-fold depletion of *tbxta* mutant cells in the presomitic mesoderm relative to the control *tyr* mutant cells (**Fig. 1F,G**). Additionally, *rx3* mutant embryos have defects in the specification of retinal neurons from early forebrain progenitors, resulting in an eyeless phenotype (*21*). Consistent with this phenotype, we observed a depletion of *rx3* mutant cells among all retinal cell types compared to *tyr* mutant cells (**Fig. S5A,B**). Therefore, MIC-Drop-seq is capable of detecting phenotypes that enrich or deplete particular cell types.

Some mutant phenotypes manifest as altered gene expression without changing the abundance of a given cell type. To determine whether MIC-Drop-seq can also detect these phenotypes, we tested for differential gene expression between mutant genotypes in each cell type. *cdx4* has a well characterized role in regulating hox family gene expression domains in the spinal cord (*22, 23*). We identified *cdx4* mutant cells in the spinal cord and compared their gene expression profile to all other cells in these clusters (**Fig. 1H**). Expression of *hoxb9a, hoxc6b*, and *hoxa9a* was significantly downregulated among *cdx4* mutant spinal cord cells (**Fig. 1I**). Taken together, these experiments demonstrate that MIC-Drop-seq is a reliable and sensitive method for multiplexed mutant phenotyping with cellular resolution.

### A 50 gene MIC-Drop-seq screen for early zebrafish development phenotypes

We next applied MIC-Drop-seq in a large-scale screen in an effort to discover novel developmental phenotypes. We selected a set of 50 transcriptional regulator genes that are variable in expression across cell types in the zebrafish embryo and highly expressed in at least one cell type (**Fig. S6A**, see Methods for details). These transcription factors are expressed across a wide variety of cell types and have roles in development that range from well-characterized to mostly uninvestigated (**Fig. S6B**).

A MIC-Drop library containing droplets targeting these 50 genes for mutagenesis was generated and injected into 1,000 zebrafish embryos (**Table S3**). To facilitate downstream statistical analysis, injected embryos were split into pools of 250 embryos, representing four biological replicates (**Fig. 2A**). Each pool was independently dissociated into single cells and sequenced at 24 hpf. After integration, filtering, and clustering, we recovered 226,492 cells distributed across 74 cell clusters (**Fig. 2B, Table S4**). Cell types were assigned with reference-based label transfer and broadly classified as either Neural, Mesoderm/Endoderm, or Non-neural ectoderm (**Fig. 2B, Table S4**). Genotypes were confidently assigned in 135,881 cells, with each of the 50 genes represented in each of the 4 replicates with an average of 2,718 cells per genotype (**Fig. S6C**).

**Fig. 2:**
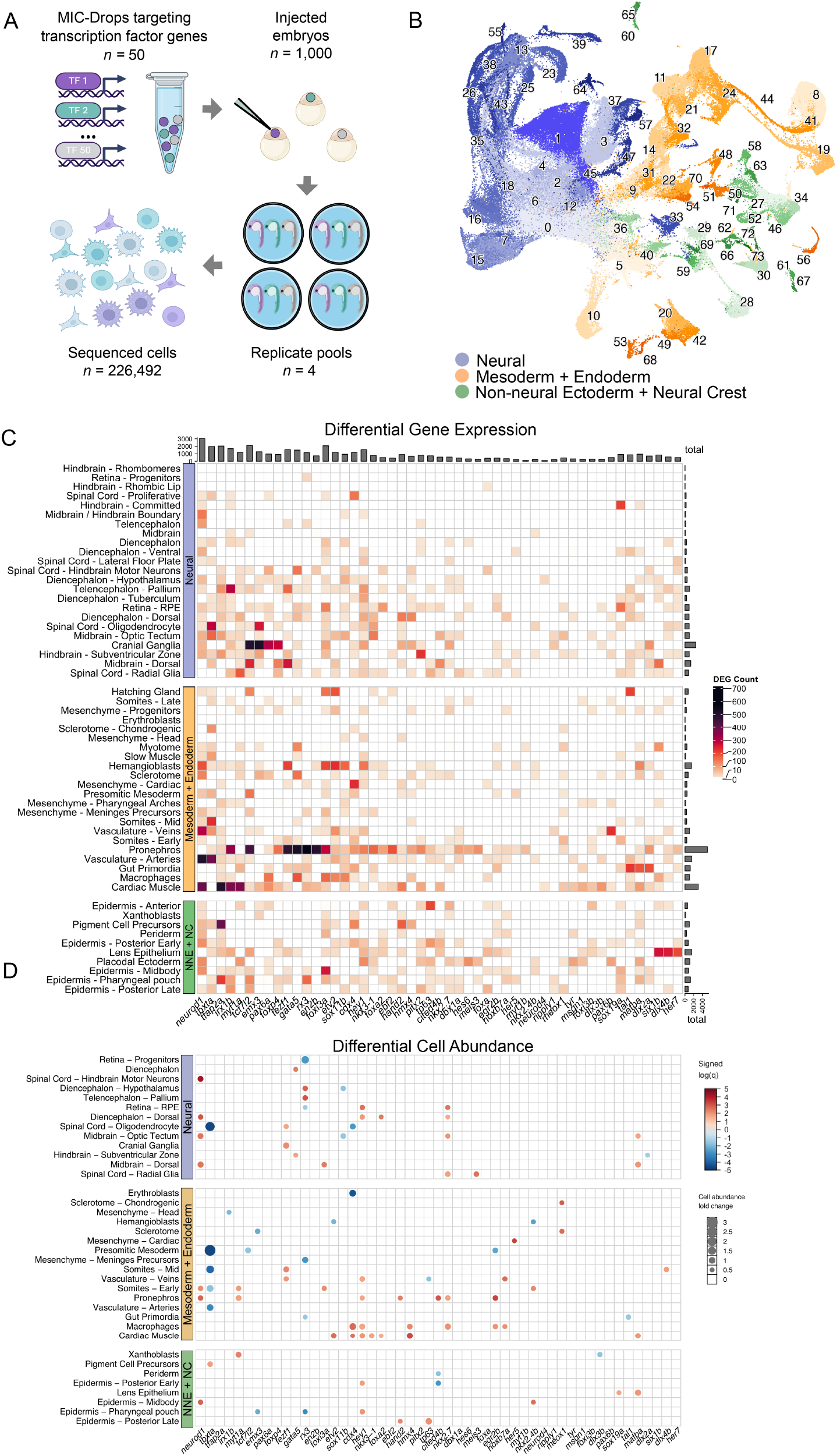
A 50 gene MIC-Drop-seq screen for early zebrafish development phenotypes. (A) Schematic of MIC-Drop-seq experiment. A MIC-Drop library was constructed with droplets targeting a set of 50 highly expressed transcription factors (see Methods) and injected into 1,000 embryos. Embryos were divided at 24 hpf into 4 replicate groups of 250 and separately dissociated into single cells for sequencing. (B) UMAP embedding of 226,492 cells derived from MIC-Drop injected zebrafish embryos. Cluster numbers indicate individual cell types; colors indicate broad tissue classifications. (C) Heatmap of differential gene expression in prominent cell types (>800 recovered cells). Color scale indicates the total number of differentially expressed genes (DEGs) in each cell type in cells with the indicated perturbation compared to all other cells in the cluster. Total DEG counts (up or downregulated) are shown for each cell type (row) and perturbation (column) (D) Bubble plot of differentially abundant cell types in prominent cell types (>800 recovered cells). Cell counts were used to calculate the fold change in cell abundance for each mutant genotype in every cell type (bubble size), while q values were calculated using monocole3 (*24*) (color scale), and signed corresponding to a depletion (- sign, blue) or enrichment (+ sign, red) for each cell type. Changes in cell abundance with q < 0.05 and absolute log2FC of > 0.5 are shown. Cell clusters with no detected cell abundance changes in any mutant were omitted.

To characterize mutant molecular phenotypes, we examined how each transcription factor disruption changed gene expression across all major cell types in the embryo. Differential gene expression testing leveraged the four biological replicates and revealed pervasive cell-type specific gene expression profiles for each mutant (**Fig. 2C**). Overall, transcription factor disruptions resulted in a median of 12 affected cell types (> 10 Differentially Expressed Genes (DEGs)) per genotype (**Fig. S7A**). Some DEGs appeared repeatedly across all perturbed cell types and mutant genotypes (**Fig. S7B**). Among these frequently observed DEGs, there was an enrichment for genes involved in Notch signaling, neurogenesis, cell cycle regulation, and stem cell differentiation, consistent with the roles of the targeted transcription factors in regulating embryonic development (**Fig. S7C**).

Mutations in developmental regulators can alter cell type differentiation trajectories and change the abundance of different tissues. We tested whether each of the 50 gene disruptions produced detectable changes in the abundance of cell types in the 24 hpf embryo. For each perturbation, fold-changes in cell type counts were calculated for each replicate, enabling significance testing of enrichment or depletion of each cell cluster (**Fig. 2D**). Overall, changes in cell type abundance were less common than transcriptional phenotypes, with a median mutant having only a single major cell type with significantly altered abundance (**Fig. S8**).

Together, these analyses demonstrate the ability of MIC-Drop-seq to comprehensively profile cellular and molecular phenotypes for hundreds of mutant animals in parallel. This rich dataset contains phenotypic reference information across hundreds of perturbation/cell type combinations. We created a web-based interactive tool to facilitate community exploration of the differential gene expression and cell type abundance data (**<u>link</u>**).

### MIC-Drop-seq enables discovery of transcriptional and cellular phenotypes

The molecular and cellular phenotypes generated from MIC-Drop-seq serve as testable hypotheses for experimental validation, enabling discovery of new biological functions of the targeted genes. Using orthogonal methods, we validated several novel phenotypes identified in our analyses. Transcriptional regulator *meox1* (mesenchyme homeobox 1) is a critical mesoderm regulator involved in somitogenesis, axial skeleton formation, and mesenchymal cell function (*25–27*). Loss-of-function mutations in *meox1* are associated with congenital scoliosis and vertebral fusions, known as Klippel-Feil syndrome, in both humans and in zebrafish (*28, 29*). The specific set of effector genes that are dysregulated in *meox1* mutants that manifest this phenotype, however, has not been determined. At 24 hpf, *meox1* is highly expressed in paraxial mesoderm cell types. Among these cells, we detected a significant reduction in expression of a set of secreted extracellular matrix proteins in *meox1* mutant cells (**Fig. 3A**). These factors, including cartilage oligomeric matrix protein (*comp*), are specifically co-expressed with *meox1* in paraxial mesoderm at 24 hpf (**Fig. 3B**), but have not previously been shown to be altered in *meox1* mutants. We injected embryos with Cas9/gRNA complexes targeting *meox1* and examined *comp* expression with in situ hybridization. In contrast to control embryos, *meox1* gRNA-injected embryos showed a loss of *comp* expression in myosepta and tailbud mesoderm, while retaining *comp* expression in the notochord (**Fig. 3C**). *Comp* has a well-documented role in matrix function, and mutations in this gene are commonly associated with disorders of the joints and skeleton in humans (*30*). Thus, our finding of dysregulated *comp* expression in *meox1* mutants demonstrates the ability of MIC-Drop-seq to uncover localized developmental phenotypes for clinically relevant genes.

**Fig. 3:**
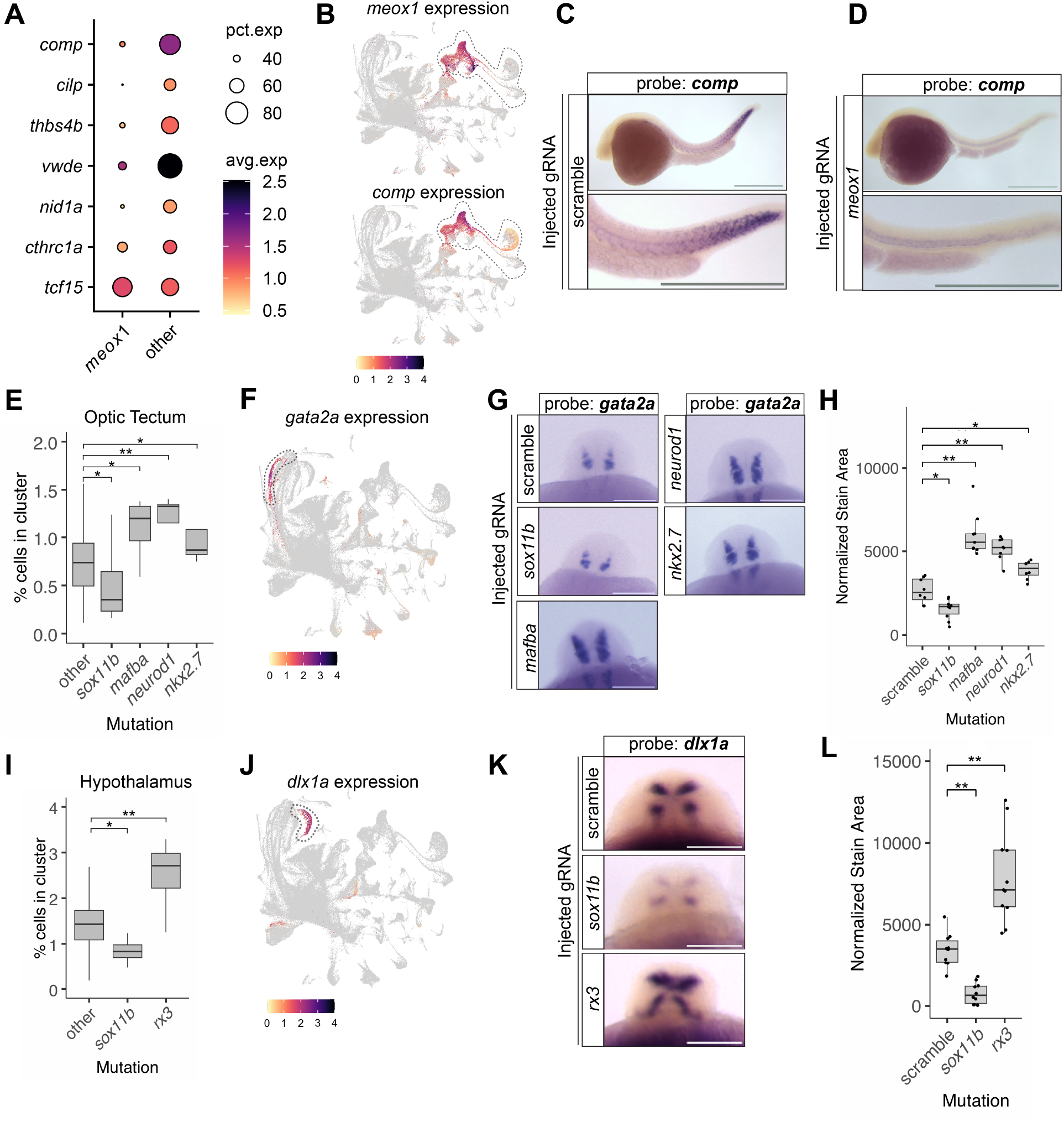
MIC-Drop-seq enables discovery of transcriptional and cellular phenotypes. (A) Gene expression bubble plot comparing expression of a set of secreted extracellular matrix factors in *meox1* mutant vs other cells in the somitic mesoderm (myotome, sclerotome, presomitic mesoderm, and early somite clusters). *tcf15* is a marker of somitic mesoderm and is unchanged in *meox1* mutant cells. Percent of cells expressing the indicated gene is indicated by bubble size and average expression level per cell is indicated with color scale. (B) Gene expression feature plots for *meox1* and *comp*. Somitic mesoderm clusters are highlighted inside the dashed line. (C-D) Colorimetric RNA in situ hybridization with DIG-labeled *comp* probe in (C) wild type and (D) *meox1* mutant 24 hpf embryos. Top: whole embryo. Bottom: zoomed in view of embryo tail. Scale bars = 400 µm. (E) Quantification of percentage of total cells of the indicated mutant in the optic tectum cluster. (F) Gene expression feature plot of *gata2a* expression in the single-cell dataset; optic tectum cluster is encircled (dashed line). (G) Representative dorsal-view images of colorimetric RNA in situ hybridization using *gata2a* probe are shown for the indicated genotype. Scale bars = 100 µm. (H) Plot quantifying *gata2a* stain area for the indicated mutants. (I) Quantification of percentage of total cells of the indicated mutant in the hypothalamus cluster. (J) Gene expression feature plot of *dlx1a* expression in the single-cell dataset, hypothalamus cluster is encircled (dashed line). (K) Representative dorsal-view images of colorimetric RNA in situ hybridization using *gata2a* probe are shown for the indicated genotype. Scale bars = 400 µm. (L) Plot quantifying *dlx1a* stain area for the indicated mutants. Significance testing in (H) and (L) was performed using unpaired t-tests comparing each mutant to scramble injected embryos. ** indicates p < 0.0005, * p < 0.005. Significance testing in (E) and (I) was performed using Hooke (part of the Monocle 3 package (*24*)). * indicates a q value of < 0.05, and ** indicates a q value of < 0.005.

Developmental phenotypes can manifest in depletion or accumulation of different cell types. From the MIC-Drop-seq dataset, several transcription factor deletions resulted in changes in neural cell type abundance in the developing brain (**Fig. 2D**). We evaluated these prospective phenotypes in the optic tectum and the hypothalamus, two cell types where multiple mutants had altered cell abundance. In the optic tectum, *mafba, neurod1*, and *nkx2*.*7* mutant cells appeared enriched while *sox11b* cells were depleted (**Fig. 3E**). We identified *gata2a* as a specific marker for optic tectum, similar to the orthologous Gata2, which marks photorecipient neurons in the mammalian tectum (*31*) (**Fig. 3F**). In situ hybridization of *gata2a* confirmed the expansion of the optic tectum in the brains of the *neurod1, mafba, nkx2*.*7* and a decrease in *sox11b* gRNA-injected embryos (**Fig. 3G,H**). We observed similar dynamic changes in cell abundance in the hypothalamus cell cluster, where eyeless *rx3* mutant cells are enriched and *sox11b* cells are depleted (**Fig. 3I**). In situ hybridization of *dlx1a*, a marker of the hypothalamus (*32*) (**Fig. 3J**), in *sox11b* gRNA-injected embryos confirmed a depletion of hypothalamic cells compared to control injected embryos, while *rx3* gRNA*-*injected embryos showed a marked increase (**Fig. 3K, L**). Critically, none of the affected mutants showed changes in expression of marker genes *gata2a* and *dlx1a* on a per-cell basis (**Fig. S9)**. Together, these data define new roles for several transcription factors in brain cell type differentiation, demonstrating how MIC-Drop-seq can uncover previously uncharacterized regulatory networks that drive organ development.

### MIC-Drop-seq reveals pervasive cell-extrinsic effects of transcription factor disruption

Cells in developing embryos can influence the behaviors of other cell types, often via direct contact or through secreted signals. These cell-extrinsic effects are often termed non-cell autonomous or indirect, and can be regulated by the cell-intrinsic activities of transcription factors. These emergent phenotypes can be absent in simplified cell culture systems and are difficult to predict in complex whole-animal systems. We reasoned that the large number of profiled genotypes in the MIC-Drop-seq dataset represents an avenue for unbiased identification and classification of these cell-extrinsic phenotypes. We define perturbation phenotypes as cell-intrinsic when the target transcription factor is expressed in the affected cell type or lineage-extrinsic when expressed in progenitors, while effects on cell types without current or prior expression of the perturbed gene are considered cell-extrinsic (**Fig. 4A**). To classify each transcriptional phenotype as cell-intrinsic or cell-extrinsic, we integrated the MIC-Drop-seq dataset with the Daniocell dataset, a time-course scRNAseq atlas of zebrafish embryogenesis (*8*). We constructed a simplified tree of cell type differentiation, using prior studies (*9, 10*), to establish the identities and trajectories from early blastula to cell types present at 24 hpf (**Fig. 4B**). This representation allows quantification of the expression of each transcription factor throughout the differentiation history of each cell type (**Fig. S10**).

**Fig. 4:**
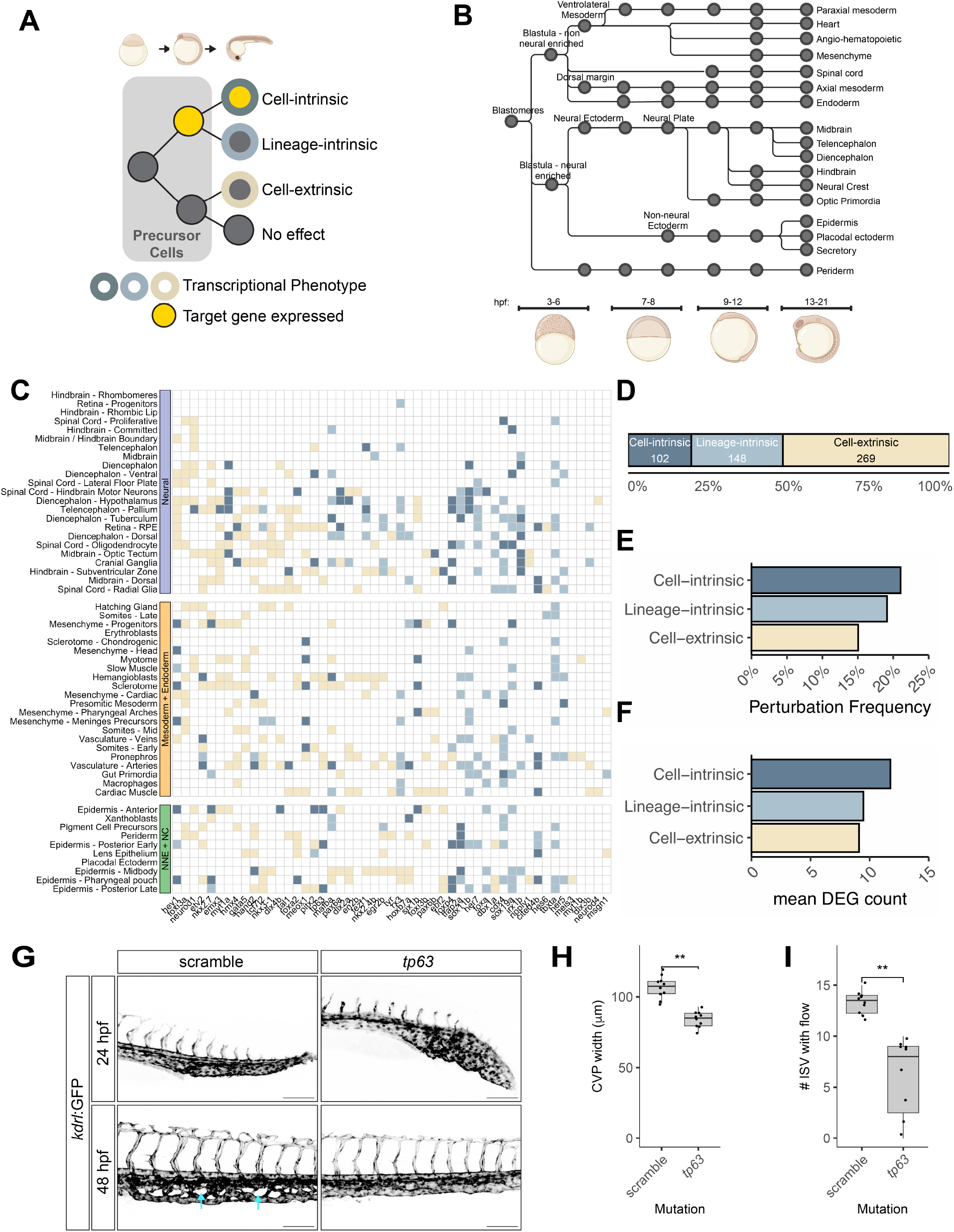
MIC-Drop-seq reveals pervasive cell-extrinsic effects of transcription factor disruption. (A) Schematic of classification of perturbation types. Perturbed cell types with significant differential gene expression can be a result of direct cellular effects (target gene currently expressed in perturbed cell type), direct lineage effects (target expressed only in precursor cells of the same lineage), or indirect effects (target not expressed in perturbed cell type or its precursors). (B) Schematic of zebrafish cell differentiation trajectories. Each cell type label is derived from the Daniocell reference dataset, with literature-based lineage connections. (C) Categorical map of perturbation effects. Detected phenotypes were classified as cell-intrinsic when the target gene is expressed in the affected cell type (dark blue) or precursor cells (light blue), while phenotypes in cell types with no history of expression were classified as cell-extrinsic (yellow). Clusters with <10 DEGs are classified as not affected (see Methods for details). (D) Proportion plot of each perturbation type within the dataset. Totals for each perturbation type are indicated. (E) Quantification of perturbation frequency for each category. For each target, expression history was determined for every cell type (cell, lineage, or indirect). The frequency of cell type perturbation (>10 DEGs) for each category is plotted from all perturbations. (F) The mean number of DEGs for perturbed cell types (>10 DEGs) for each category is plotted. (G) Representative greyscale fluorescence images of CVP and posterior trunk of *kdrl*:GFP labeled transgenic embryos injected with scrambled or *tp63* targeting gRNA at 24 and 48 hpf. Blue arrows highlight CVP fenestrations in control embryos. Scale bar = 100 µm. (H) Measured CVP pixel width at 48 hpf in injected embryos. (I) Quantification of ISVs with detectable erythrocyte flow.

We next quantified the cell-intrinsic and cell-extrinsic impacts of each transcription factor disruption using lineage and target gene expression relationships. We set a simple threshold of > 10 DEGs to define a transcriptional phenotype and computed the scaled expression of each target gene across each cell type. Phenotypes were classified as cell-intrinsic when the affected cell types or their precursors expressed the target gene or were otherwise classified as cell-extrinsic (**Fig. 4C**). Overall, we found that cell-extrinsic effects are pervasive, representing the most commonly observed phenotype class among all affected cell types (**Fig. 4D**). Although cell-extrinsic effects were the most common class overall, individual cell types showed stronger predisposition to transcriptional changes when the target gene was expressed in those cells or their precursors **(Fig. 4E)**. Moreover, these cell-intrinsic phenotypes tended to have larger transcriptional changes, reflected by a higher average number of DEGs **(Fig. 4F)**.

To verify the veracity of cell-extrinsic phenotypes observed by MIC-Drop-seq, we focused on vascular cell types, which displayed cell-extrinsic phenotypes in multiple mutants. For example, disrupting *tp63*, a transcription factor critical for epidermal cell development (*33*), caused a decrease in venous cell abundance (**Fig. S11A**) and altered gene expression in all vascular cell types, indicating atypical vascular development (**Fig. S11B**). These phenotypes are unexpected, given that *tp63* is not expressed in vascular cells but is instead restricted to the epidermis (**Fig. S11C**). To verify dysregulated vascular development in *tp63* mutants, a transgenic *kdrl:GFP* zebrafish line was used to compare vascular patterning in embryos injected with either *tp63-*targeting or scrambled gRNAs. At 24 hpf, *tp63* mutants displayed altered caudal vein plexus (CVP) morphology (**Fig. 4G**). At 48 hpf, vascular development remained disrupted, with failure to form typical vascular fenestrations in the CVP of *tp63* mutants (**Fig. 4G**) and displayed decreased CVP width (**Fig. 4F**). Notably, length of intersegmental vessels (ISVs) remained unchanged (**Fig. S11D**). Optical flow analysis revealed defects in erythrocyte transit in *tp63* mutants (**Fig. S11E**), including a significant decrease in the number of ISVs with detectable flow (**Fig. 4I**), and overall decreased flow area (**Fig. S11F**). Intussusceptive angiogenesis in the CVP is linked to shear force supplied from cardiac output (*34*), and p63 null mice have defects in heart formation (*35*). While *tp63* injected zebrafish embryos had defects in skin formation (**Fig. S11G**), no changes in heart rate were apparent (**Fig S11H**). We hypothesize that these defects in early vasculogenesis could be the result of altered physical forces stemming from an improperly formed epidermal layer. Together, these findings demonstrate how the unbiased whole-animal datasets generated by MIC-Drop-seq can detect unanticipated emergent phenotypes, even in cell types where the target gene is not expressed.

Profiling mutant phenotypes with scRNAseq provides a way to comprehensively map the link between genotype and the networks of genes, cells, tissues, and organs underlying animal development. In this report, we present MIC-Drop-seq as a technique for pooled CRISPR screening in whole zebrafish embryos with cellular resolution phenotyping across all embryonic cell types. MIC-Drop-seq is related to existing multiplexed scRNA-seq technologies like CROP-seq and Perturb-seq, but offers several differentiating features that enable its use in whole animals. CROP-seq and Perturb-seq similarly rely on gRNA sequencing to index mutant genotypes, but have been limited to screens of cultured cells, limiting the scope of observable phenotypes to a subset of molecular networks active in a handful of cell types (*15, 36–38*). *In-vivo* Perturb-seq uses transduction of CRISPR-containing viral particles to create clones with differing mutant genotypes within individual animals (*18, 39*). This approach, while useful for studying tissue-specific regulatory networks, fails to capture earlier developmental phenotypes of the targeted genes or to isolate gene-specific cell-extrinsic effects. MIC-Drop-seq uses gene disruption at the zygotic stage, where mutations are rapidly generated and inherited by early progenitors during development, enabling whole-animal developmental phenotypes to be profiled with cellular resolution at scale. Furthermore, MIC-Drop-seq uses injection of purified Cas9 RNP complexes instead of viral vectors, dramatically reducing the possibility of genotyping errors from ambient gRNA or cells infected with multiple viral vectors. These features make MIC-Drop-seq a highly efficient platform that could be scaled to study hundreds to thousands of whole-animal mutants in parallel.

The cellular-level comparisons made possible by MIC-Drop-seq enable detection of phenotypes that would otherwise be difficult to detect by conventional observation methods or would not manifest *in vitro*. For example, we identified and validated widespread cell abundance changes in the developing brain of *sox11b* mutant embryos, which appear visually indistinguishable from wildtype siblings and have been classified as developmentally normal in previous studies (*40, 41*). Furthermore, our transcriptomic profiling revealed extensive changes in the makeup of the myosepta extracellular matrix in *meox1* mutants, despite only having a visible phenotype at much later stages of development. While recent studies have also leveraged parallel scRNAseq of multiple mutant embryos to detect such cryptic developmental phenotypes (*42*), MIC-Drop-seq drastically increases throughput by enabling rapid delivery and demultiplexing of perturbations. Indeed, our approach enabled the simultaneous collection of phenotypic information for 50 mutants across 74 embryonic cell types in a single experiment carried out in a single day. This unbiased approach revealed pervasive cell-extrinsic phenotypes, including an unexpected emergent phenotype in the vasculature of embryos with disrupted *tp63*, a transcription factor that is expressed only in ectoderm in the 24 hpf embryo.

Our MIC-Drop-seq experiments focused on the 24 hpf zebrafish embryo, but further modifications might enable a broader set of applications. MIC-Drop-seq is in principle compatible with any species where embryos are amenable to injection at the zygotic stage. While we demonstrate efficient gRNA sequence recovery at 24 hpf zebrafish embryos, studying later developmental time points will likely result in lower recovery as the gRNAs are diluted and degraded. DNA barcodes or chemically-stabilized gRNAs could enhance genotype recovery at later stages (*43*). Supplementary oligonucleotide-based indexing of individual embryos could further aid in profiling more advanced developmental stages through genotype imputation from cells where gRNAs are not recovered (*42*). Another challenge of whole-animal scRNAseq as a phenotyping method is the detection of transcriptional effects in low-abundance cell types. More statistical power could be acquired by leveraging emerging scRNAseq technologies to sequence far more cells across more replicates (*44, 45*). Alternatively, cell sorting could enrich rare cell types of interest for focused screens.

MIC-Drop-seq is a flexible phenotyping platform that can be integrated with other perturbation types or single-cell methods in the future. For example, combining MIC-Drop-seq with scATAC or scCUT&TAG (*46*) would enable deeper probing of developmental gene regulatory networks. MIC-Drop-seq could be used to deliver and characterize the combinatorial effects of chemical and genetic perturbations in pharmacogenetic screens. More precise genome engineering methods, like base or prime editing (*47*), could enable the simultaneous profiling of phenotypes of panels of specific genetic variants. Finally, MIC-Drop-seq could be coupled with other quantitative and high-throughput measures of phenotype, including body morphology, neural activity, and behavior. Phenotypic data from systematic perturbation screens like these can be used to train predictive models of animal development (*48–50*). Large-scale molecular phenotyping of mutant animals with methods like MIC-Drop-seq will be crucial to providing training data to bridge the gap between molecular and higher-order organismal phenotypes. Such models will aid in connecting, interpreting, and predicting the gene regulatory networks that build a functional animal.

## Acknowledgments

We thank all members of the Gagnon, Parvez and Peterson labs for discussions and comments. We thank Opal Allen, Jeffrey Farrell, Austin Green, Nathan Lord, Aaron McKenna, Tajeshree Prakash, Abhinav Sur, Lisandra Vila Ellis, Yogesh Goyal and Michael Werner for feedback on the manuscript, protocols, reagents, and equipment. We thank CZAR and CBRZ staff for zebrafish care. We thank the Cell Imaging and High-Throughput Genomics core facilities.

## Funding

National Institutes of Health grant F32HL156644 (CMC)

National Institutes of Health grant K01HG013682 (CMC)

National Institutes of Health grant R00HG012593 (SP)

National Institutes of Health grant R01GM134069 (RTP)

National Institutes of Health grant R35GM142950 (JAG)

National Institutes of Health grant R24OD035409 (RTP, JAG)

Northwestern University Startup Funds (SP)

One Utah Data Science Hub (JAG)

## Author contributions

Conceptualization: SP, RTP, JAG, CMC, ZJB Methodology: SP, CMC, ZJB

Investigation: SP, CMC, ZJB, BWB, CJY Visualization: CMC, SP, ZJB

Funding acquisition: RTP, JAG, SP, CMC, ZJB Project administration: RTP, JAG

Supervision: RTP, JAG

Writing – original draft: CMC, SP, ZJB

Writing – review & editing: RTP, JAG, CMC, SP, ZJB, BWB

### Data and materials availability

All raw sequencing data, cell by gene expression matrices, and processed Seurat objects will be available on SRA.

All code and supporting data used to process data and generate visualizations in the manuscript are available at: https://github.com/clay-carey/MIC-Drop-seq

## Supplementary Materials

Materials and Methods

Figs. S1 to S11

Tables S1 to S4

References (*51-60*)

