## Supplemental Figures for "MIC-Drop-seq: Scalable single-cell phenotyping of mutant vertebrate embryos"

Figures S1-S11

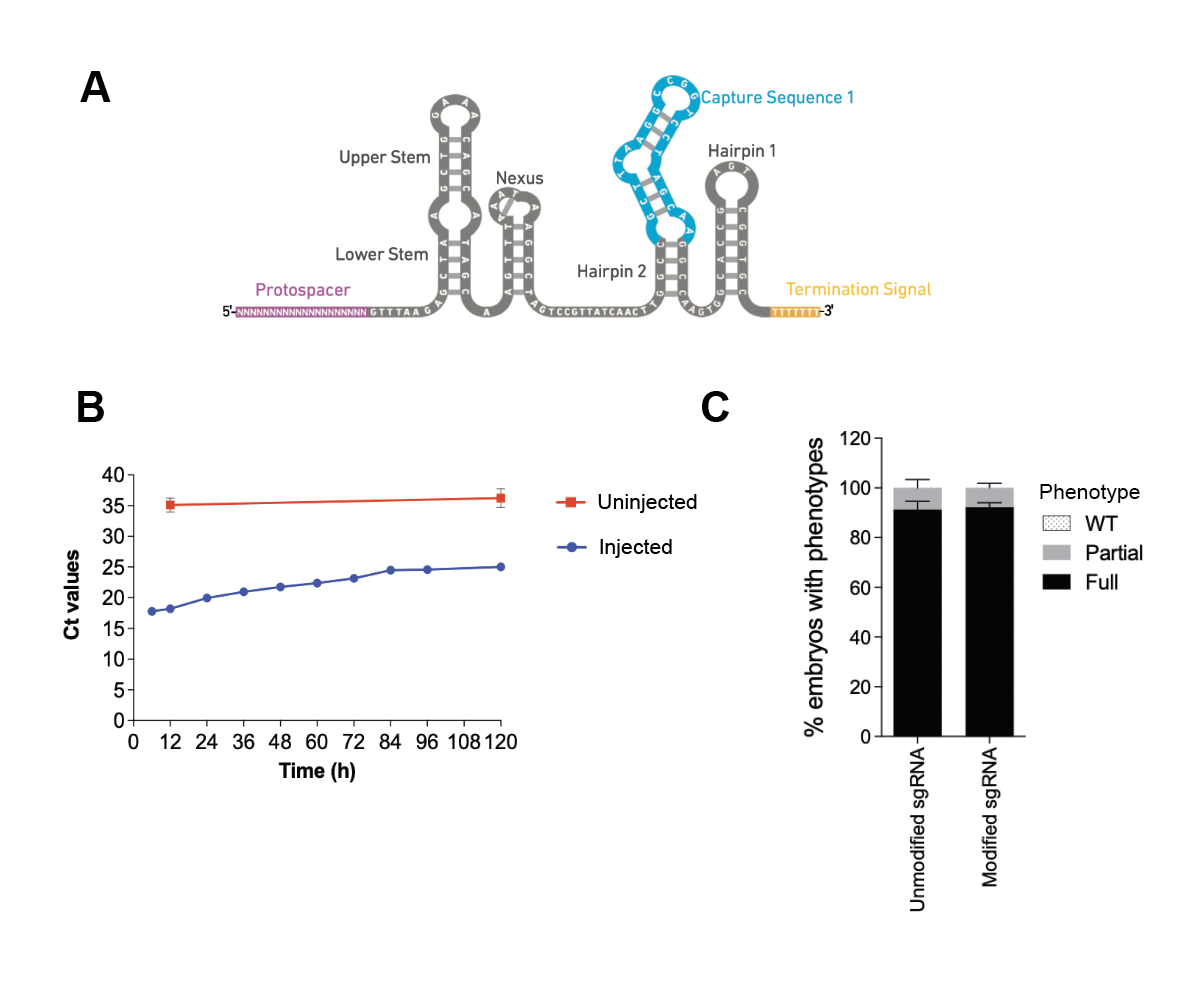

Fig S1: Modified gRNA scaffolds enable capture without impacting function.
(A) Schematic of the modified gRNA used in this study, with the capture sequence shown in blue. Image courtesy of 10X Genomics. (B) Quantitative RT-PCR Ct values for amplifying modified gRNAs from zebrafish embryos at the indicated time point after injection of purified gRNA (blue) compared to uninjected controls (red).( C) Phenotypic penetrance of embryos injected with Cas9 protein and modified or unmodified gRNA targeting the *rx3* gene, which is required for eye formation. Phenotypes were categorized as full (eyeless), partial (small eyes or one eye), or WT (no phenotype).

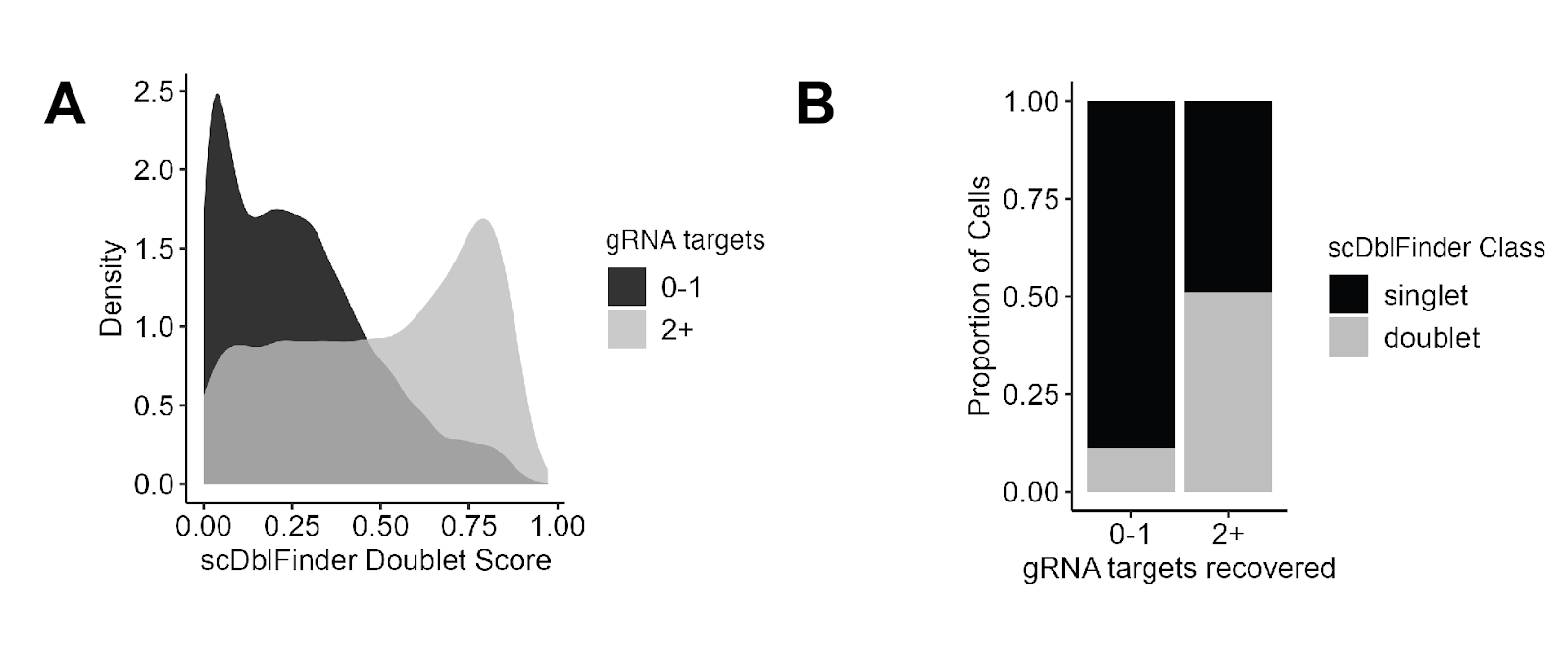

**Figure S2: MIC-Drop-seq enables multiplexed mutant phenotyping by scRNAseq.**
(A) Probability densities of scDoubletFinder scores for cells classified as having a single gRNA target (black) and cells with multiple gRNA targeting multiple genes (grey). Higher scores indicate a higher probability of a cell doublet. (B) Proportion of cells classified by scDoubletfinder as a singlet or doublet in cells with gRNA targeting 0-1 or 2+ genes.

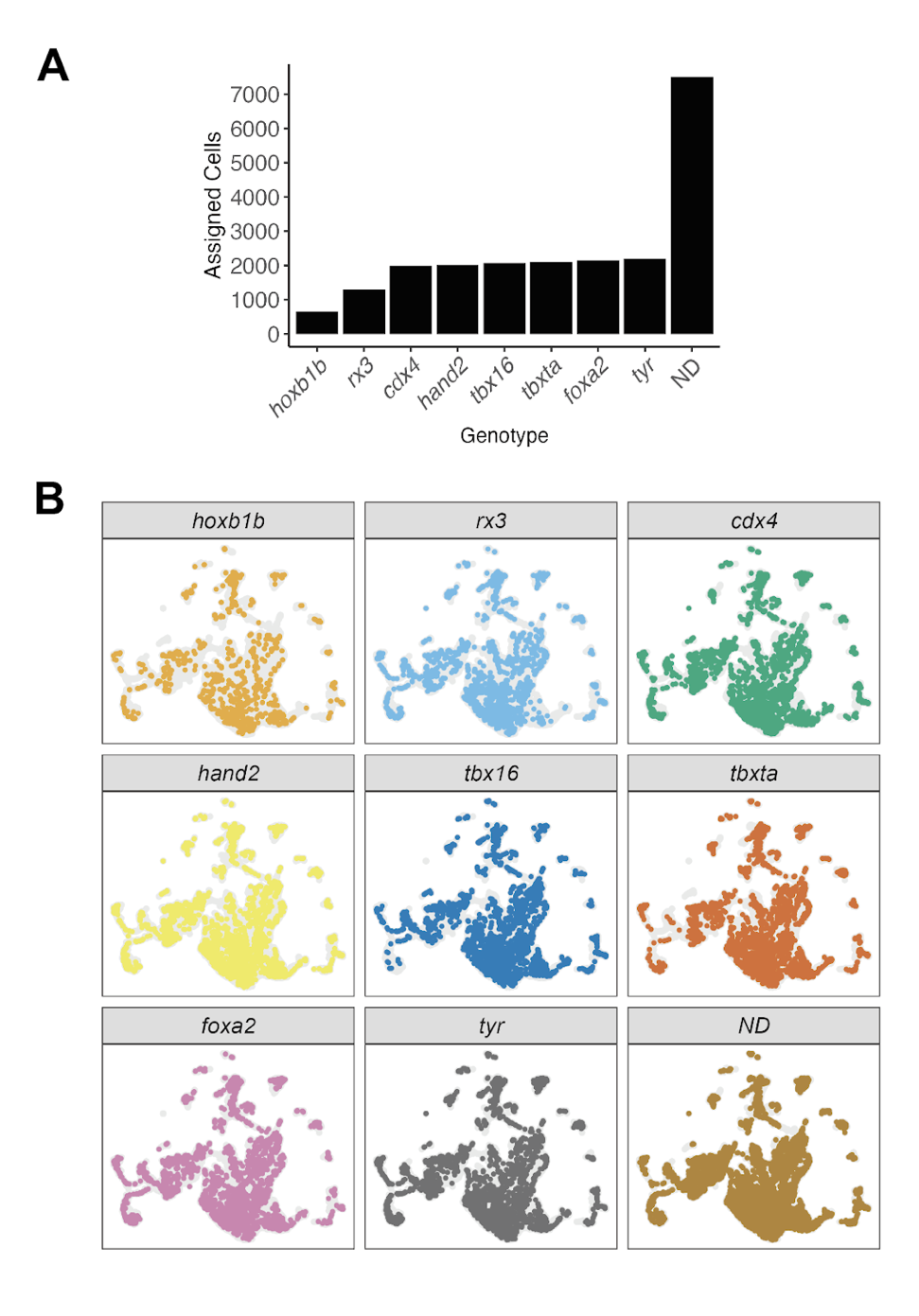

**Fig. S3: Classifying and demultiplexing mutant genotypes.**
(A) Counts of cells classified for each mutant genotype. ND = No genotype determined. (D) UMAP of dataset split and colored by predicted mutant genotype.

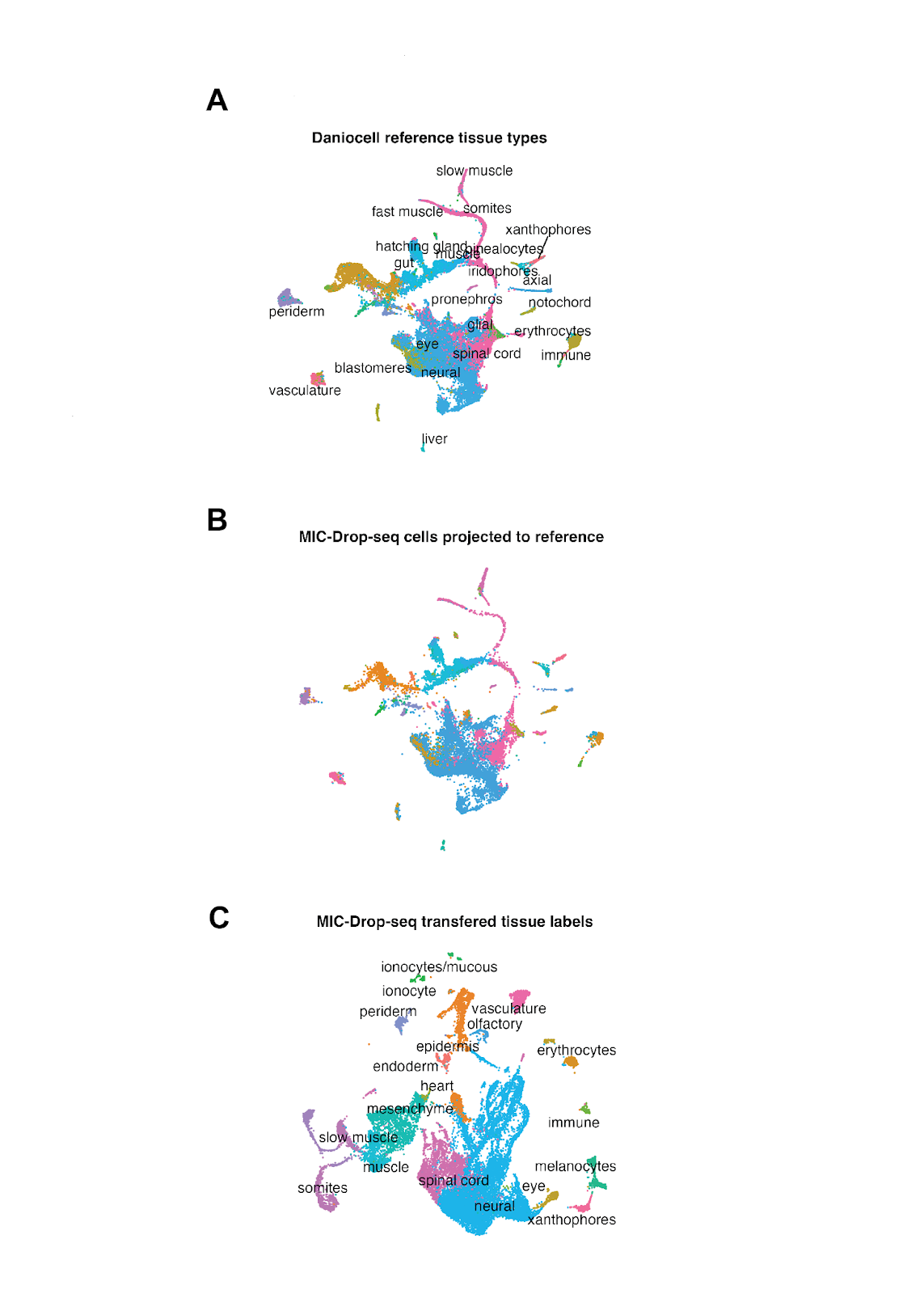

**Fig. S4:** **Reference-based label transfer.**
(A) UMAP embedding of Daniocell dataset cells from the 22-26 hpf timepoints. (B) MIC-Drop-seq experiment cells projected to the reference UMAP space. (C) MIC-Drop-seq cells embedded in native UMAP space with transferred labels.

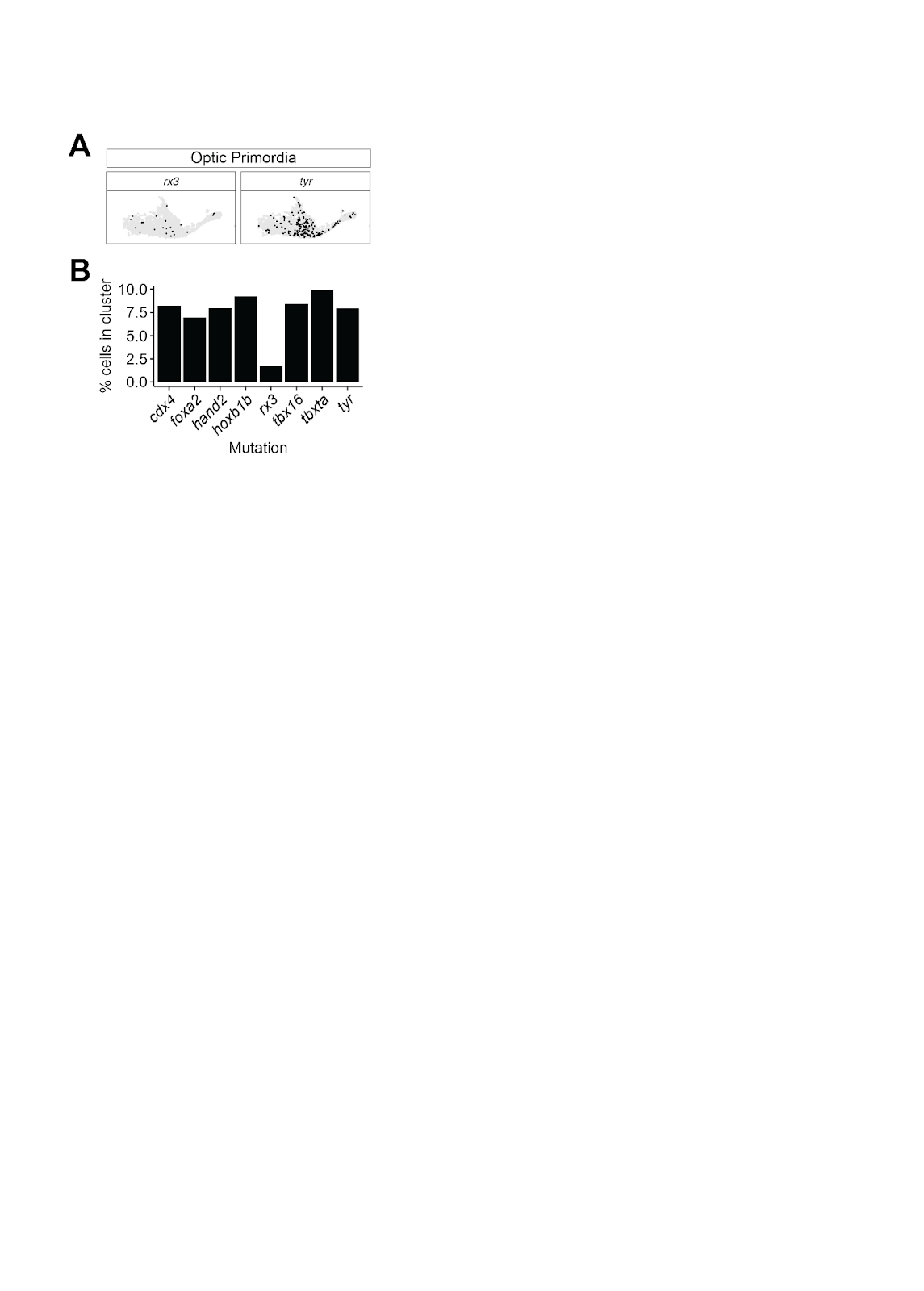

Fig. S5: Depletion of optic primordia cells in *rx3* mutants.
(A) Separate UMAP embeddings of optic primordia cells (retinal progenitors and retinal pigmented epithelium) highlighting cells with the indicated genotype. (B) Quantification of the percentage of cells in the optic primordia clusters for each indicated genotype.

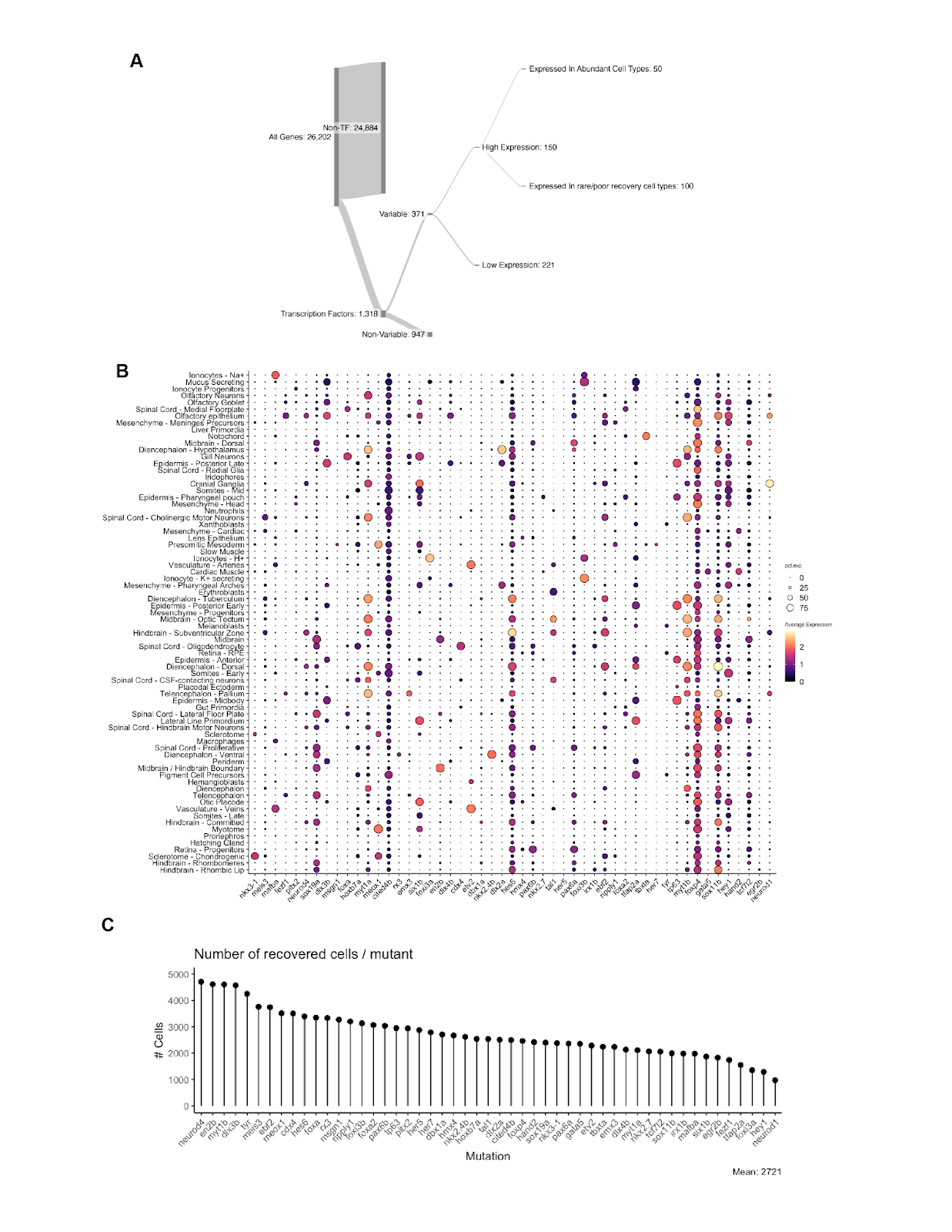

**Figure S6: Design and summary of 50 gene MIC-Drop-seq screen.**
(A) A sankey diagram with selection criteria is shown. Transcription factors with variable expression across cell types were identified. The 50 transcription factors with the highest expression in abundant cell types were selected for the MIC-Drop-seq screen. (B) Gene expression bubble plot for each transcription factor targeted in the experiment in each 24 hpf zebrafish cell type. (B) Total cell counts assigned with the indicated mutant genotype across the dataset.

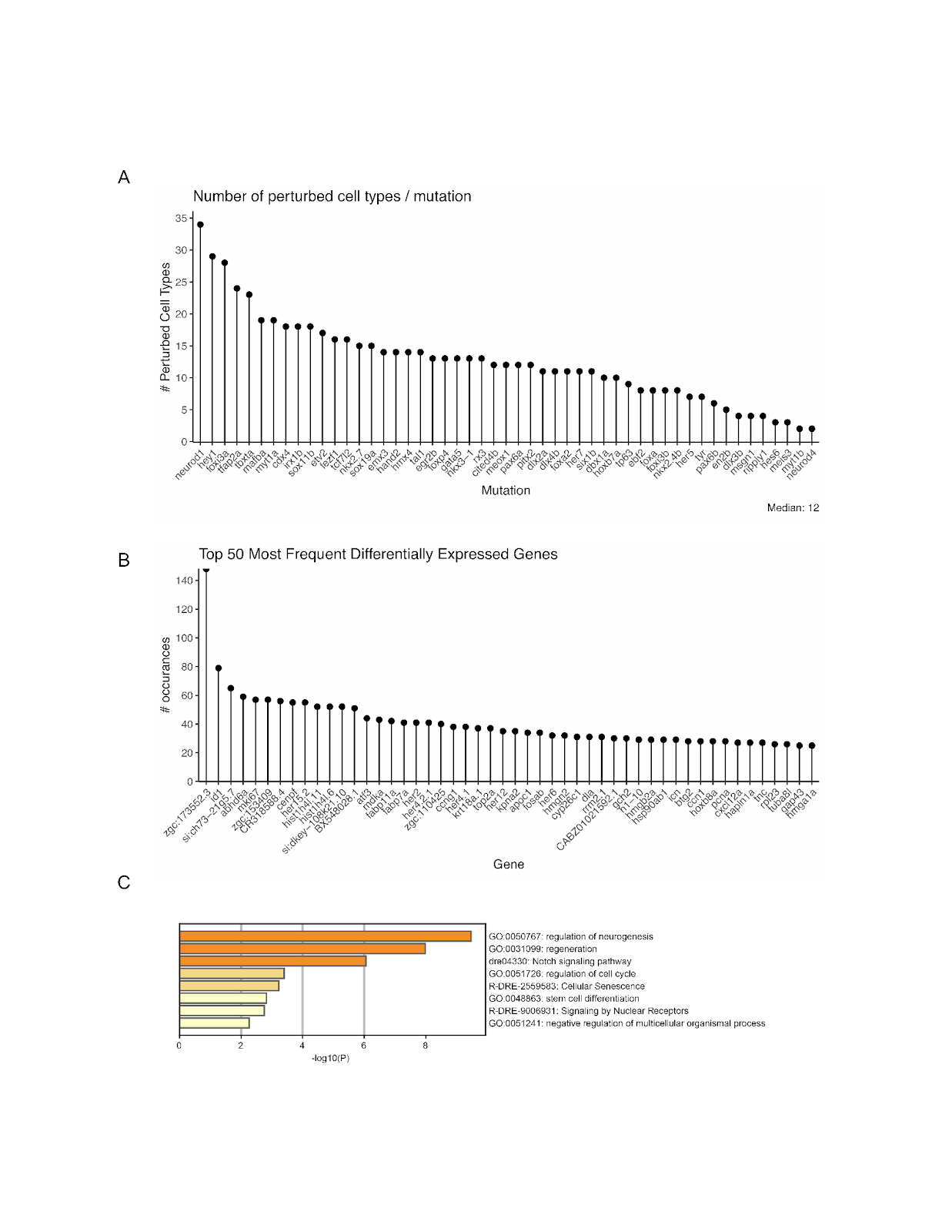

Fig. S7: Overview of transcriptional phenotypes.
(A) Quantification of the total number of significantly perturbed abundant cell types (>800 recovered cells, >10 DEGs) for each mutant. (B) Counts of the most frequently differentially expressed genes across all mutants and cell types. (C) Gene Ontology analysis of the set of the top 50 most frequently differentially expressed genes using Metascape (*60*).

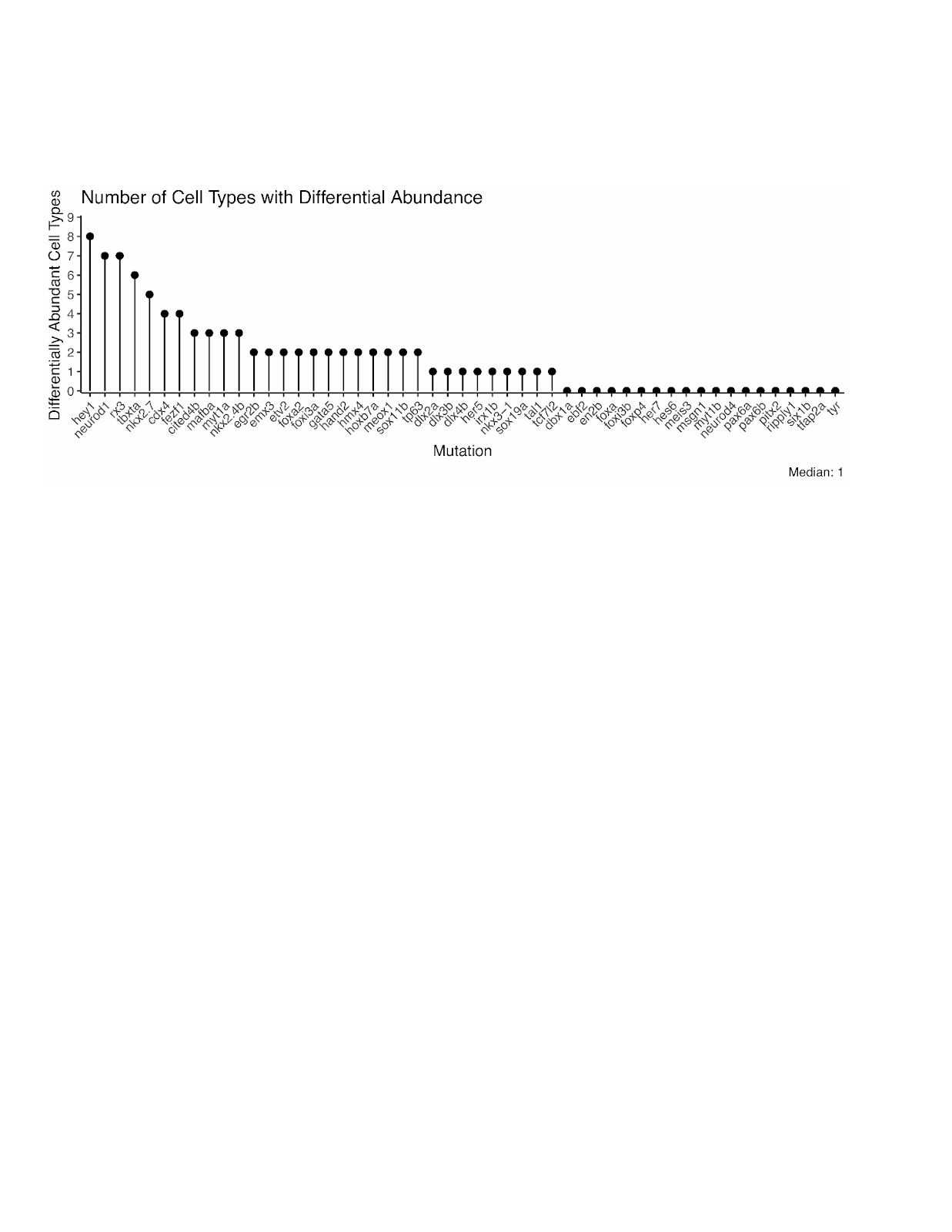

Fig. S8: Overview of cell abundance changes from MIC-Drop-seq screen.
Quantification of cell types with significantly differentially abundance for each mutant (log_2_FC > 0.5, Hooke FDR < 0.05)

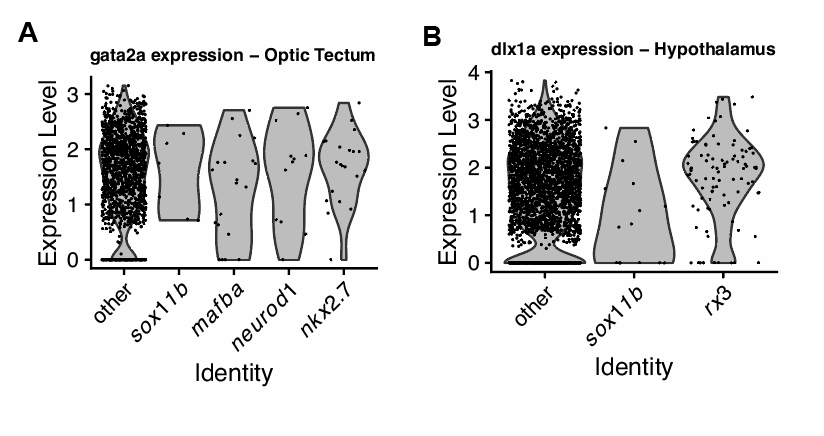

Fig. S9: Cluster marker gene expression.
(A,B) Gene expression violin plots for cells with the indicated genotype for (A) *gata2a* expression in the optic tectum cluster or (B) *dlx1a* in the hypothalamus cluster.

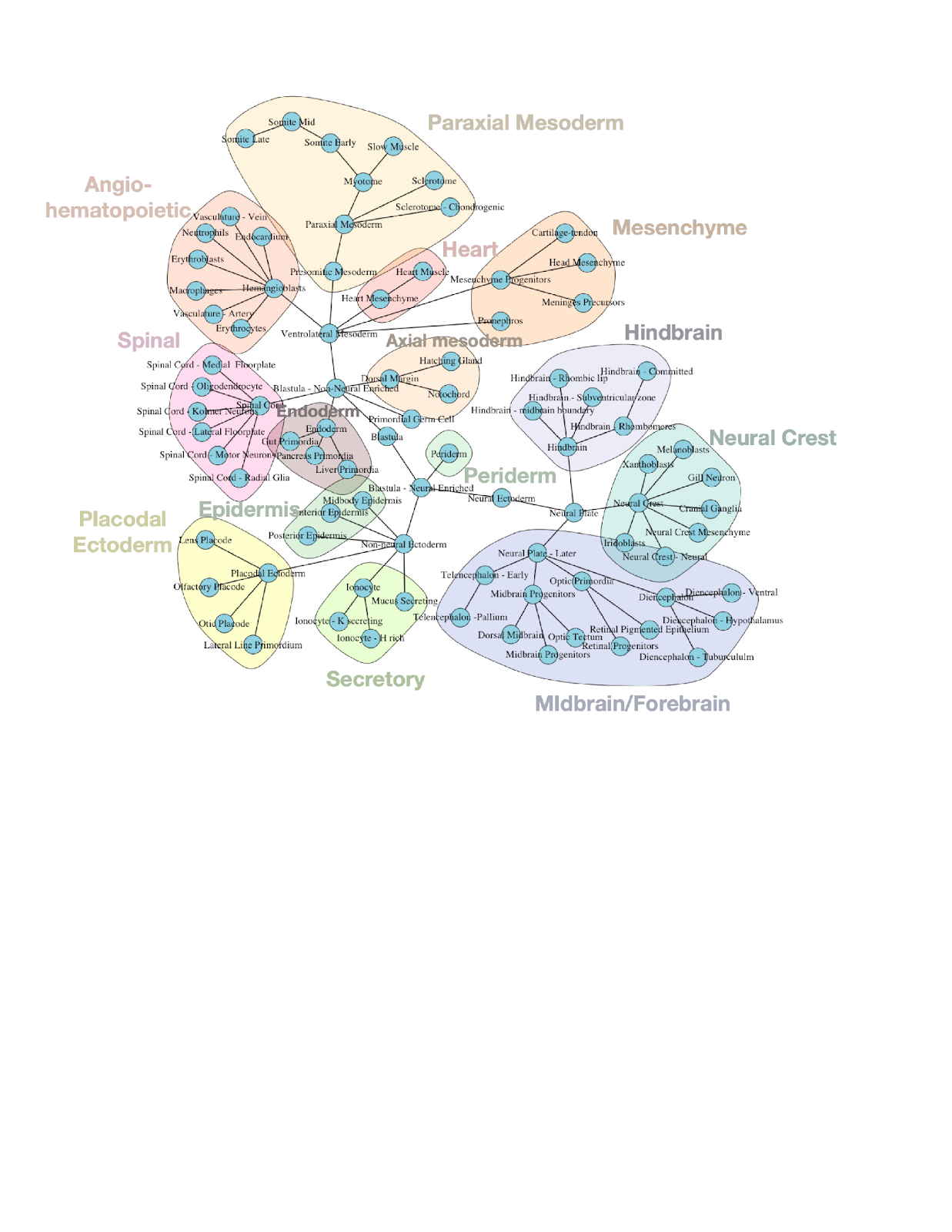

**Fig.S10:** **Differentiation schematic of 24 hpf zebrafish cell types.**
Full differentiation network map connecting cell types present in 3-24 hpf Daniocell data and MIC-Drop-seq data at 24 hpf.

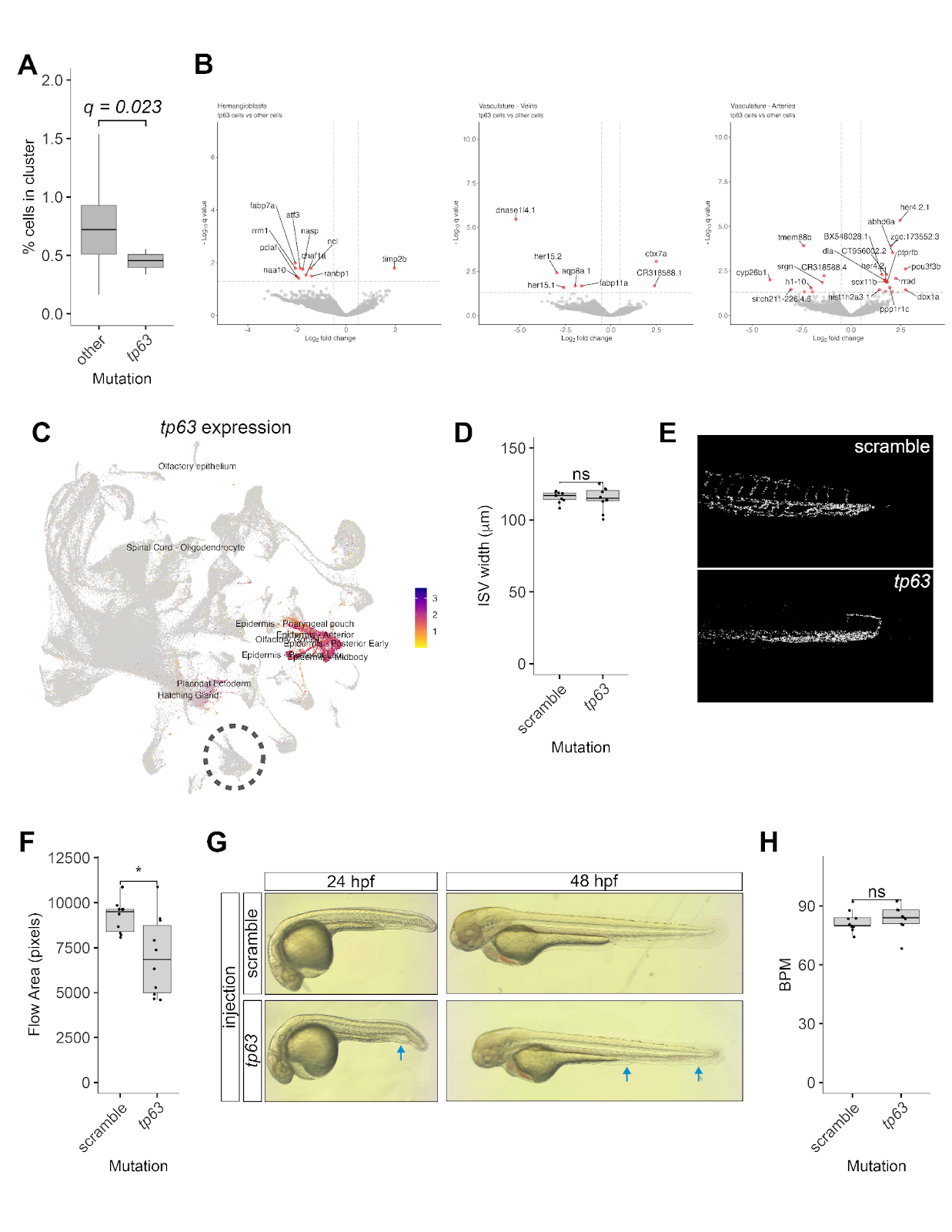

Fig. S11: Characterizing a cell-extrinsic vascular phenotype in *tp63* mutants.
(A) Quantification of venous cell percentage in *tp63* mutant vs all other genotypes from the MIC-Drop-seq dataset. Hooke FDR value is shown. (B) Volcano plots of differential gene expression in *tp63* mutant cells vs all other genotypes in hemangioblasts (left), veins (center), and arteries (right). (C) Feature plot of *tp63* gene expression. Cell types in the top 20% of scaled *tp63* expression are labeled. Vascular cells are encircled (dashed line) and do not express *tp63*. (D) Quantification of ISV width in injected 48 hpf embryos. (E) Summary images of erythrocyte flow analysis in 48 hpf injected embryos, flow is registered as white pixels. (F) Quantification of total pixel flow area from flow analysis. (G) Brightfield images of *tp63* and scramble injected embryos at 24 and 48 hpf, blue arrows indicate areas of altered epidermal development in *tp63* mutants. (H) Quantification of heart beats per minute (BPM) in scramble and *tp63* injected embryos at 48 hpf. Significance testing (D),(F), and (H) was performed using unpaired t-tests comparing each mutant to scramble injected embryos. * indicates  p < 0.05.
